# Unveiling the Role of PHF20 in TBC1D4-Mediated Glucose Uptake During *Toxoplasma gondii* Infection

**DOI:** 10.1101/2025.08.08.669278

**Authors:** Fei-Fei Gao, Xin-Cheng Wang, Guan-Hao Hong, Yu-Sun Yun, In-Wook Choi, Jaemin Yuk, Wei Zhou, Xintian Chen, Jongsun Park, Guang-Ho Cha

**Author notes:** Address correspondence and reprint requests to Dr. Xintian Chen, Laboratory of Gastroenterology, Affiliated Hospital of Guangdong Medical University, Zhanjiang 524001, Guangdong, China.; Email address and Dr. Jongsum Park, Department of Pharmacology, Chungnam National University, School of Medicine, Daejeon 35015, Republic of Korea.; E-mail address, and Dr. Guang-Ho Cha Department of Infection Biology, Chungnam National University, School of Medicine, Munhwa-dong, Jung-gu, Daejeon 35015, Republic of Korea. These authors contributed equally to this work (Fei-Fei Gao, Xin-Cheng Wang).

## Abstract

**Background:** *Toxoplasma gondii* relies on host cell nutrients, especially glucose, to support its intracellular growth. However, the mechanisms by which it enhances host glucose metabolism remain incompletely understood.

**Methods:** We used fluorescence-based glucose uptake assays, flow cytometry, metabolic flux analysis, and gene silencing in ARPE-19 cells, along with transgenic mouse models, to explore how *T. gondii* manipulates host glucose utilization and whether this promotes parasite proliferation.

**Results:** *T. gondii* infection significantly enhanced host glucose uptake in a dose-dependent manner and promoted GLUT4 translocation to the plasma membrane. Notably, extracellular acidification rate (ECAR) assays demonstrated a marked increase in glycolytic activity following infection. Mechanistically, we identified that the PI3K/AKT signaling pathway mediates the phosphorylation and transcriptional upregulation of TBC1D4, a Rab-GAP protein essential for GLUT4 trafficking. Genetic silencing of TBC1D4 impaired both glucose uptake and parasite replication. Furthermore, we uncovered a regulatory mechanism involving AKT-dependent nuclear signaling that modulates TBC1D4 transcription. In vivo experiments using Phf20 transgenic mice confirmed increased susceptibility to *T. gondii* and elevated glucose metabolic responses.

**Conclusions:** Our study reveals that *T. gondii* hijacks a host PI3K/AKT-TBC1D4 axis to enhance glucose uptake and glycolysis, thereby ensuring metabolic support for its proliferation. These findings highlight host glucose metabolism as a potential therapeutic target for controlling toxoplasmosis.

## Introduction

*Toxoplasma gondii* (*T. gondii*) is a ubiquitous intracellular parasite capable of infecting nearly all warm-blooded animals, representing one of the most prevalent parasitic infections globally^[1]^. Although generally asymptomatic, infections in prenatal and immunocompromised individuals can lead to severe disease, neurological disorders, and even death^[2]^. Congenital toxoplasmosis, as a result of vertical transmission from infected mothers, is a significant cause of morbidity and mortality in fetuses, neonates, and children as they progress into adulthood^[3]^. The disease produces a classical triad of hydrocephalus, intracranial calcification and chorioretinitis, namely ocular toxoplasmosis^[4]^. Yet, the indications for treatment still remain debated, and people are trying to find a way to prevent it.

To thrive and replicate successfully, *T. gondii* has evolved sophisticated strategies to exploit host nutrients, particularly glucose, to fuel its metabolic processes^[5]^. The parasite reprograms host cell metabolism to meet its high energy and biosynthetic demands, ensuring sufficient resources for its survival and replication^[6, 7]^. However, the molecular mechanisms driving this metabolic reprogramming remain poorly understood. Recent studies have revealed that *T. gondii* infection enhances glucose uptake in host cells, underscoring the critical role of glucose in parasite metabolism^[8]^. Autofluorescence lifetime imaging indicated that, over time, infected host cells showed decreases in levels of intracellular glucose and lactate^[9]^. Glucose is essential for intracellular pathogens like *T. gondii*, as it supports energy generation via glycolysis and facilitates glycoconjugate synthesis, which is vital for the parasite’s proliferation^[8, 10]^. The PI3K/AKT signaling pathway, known for regulating glucose transporter translocation, emerges as a potential route through which *T. gondii* might promote host cell glucose uptake ^[11, 12]^. In the past two decades, accumulated evidences showed that *T. gondii* infection activates PI3K/AKT signaling pathway^[13–15]^. In mammalian cells, PI3K/AKT activation leads to the phosphorylation of TBC1D4, a critical mediator of GLUT4 vesicle translocation to the plasma membrane in response to insulin signaling^[16, 17]^. However, whether *T. gondii* leverages this pathway to increase glucose availability in host cells has yet to be established.

TBC1D4 (TBC1 domain family member 4), also known as AS160, is a Rab GTPase-activating protein (Rab-GAP) that plays a pivotal role in glucose transporter trafficking^[18, 19]^. It acts as a key regulator of GLUT4 translocation in response to insulin signaling via phosphorylation by PI3K/AKT signaling pathway. TBC1D4 phosphorylation inhibits its GAP activity, enabling the release of GLUT4-containing vesicles to the plasma membrane, which enhances glucose uptake^[20, 21]^. In the context of *T. gondii* infection, TBC1D4’s regulation may provide a mechanism through which the parasite increases host cell glucose availability to meet its metabolic demands. It has been reported that *T. gondii* infection of mouse macrophages activates PKB/AKT *in vivo* and *in vitro*^[13]^, our previous data also showed that *T. gondii* infection can downregulate FAF1 in PI3K/AKT/FOXO1-dependent manner, which is essential for the parasite proliferation^[15]^. Hence, we hypothesize that *T. gondii* subverts the host PI3K/AKT-TBC1D4 signaling axis to enhance GLUT4 trafficking, facilitating glucose uptake for its survival and proliferation. Our findings demonstrate that *T. gondii* infection upregulates both the phosphorylation and transcriptional expression of TBC1D4, with both processes being mediated by PI3K/AKT activity. Additionally, we identify PHF20, a transcription factor modulated by PI3K/AKT signaling, as a novel regulator of TBC1D4 expression during infection. PHF20 overexpression or knockdown significantly affects TBC1D4 protein levels, consequently influencing glucose availability to the parasite. Our study establishes PHF20 as a pivotal regulator of TBC1D4 transcription during *T. gondii* infection, expanding its functional scope beyond oncology. By modulating GLUT4 trafficking and glucose uptake, PHF20 ensures the metabolic flexibility required for parasite survival and proliferation, marking a significant advancement in understanding the molecular basis of host-pathogen interactions.

Together, these findings elucidate a regulatory link between PHF20 and the TBC1D4/GLUT4 pathway, hijacked by *T. gondii* to meet its metabolic demands. This study provides new insights into the molecular basis of *T. gondii*-induced host metabolic manipulation, offering potential therapeutic avenues for controlling toxoplasmosis by targeting metabolic pathways critical to the parasite’s life cycle.

## Materials and Methods

### Antibodies and Reagents

The following antibodies were used: anti-TBC1D4 (ab189890) was purchased from Abcam (Cambridge, UK). Anti-GLUT4 (SC-53566), anti-α-Tubulin (SC-32293), and anti-TP3 (sc-52255) were obtained from Santa Cruz Biotechnology (Santa Cruz, CA, USA). Anti-phosphor (S/T)-AKT substrate (#9611), anti-GLUT4 (#2213), anti-phospho-AKT (Ser473) (#9271), anti-phospho-AKT (Thr308) (#13038), anti-AKT (#9272), anti-EGFR (#2232S), anti-phospho-TBC1D4 (#8881) and anti-PHF20 (#3934) were obtained from Cell Signaling Technology (Burlingame, CA, USA). Anti-LC3A-I/II (NB100-2331) was purchased from Novus Biologicals (Centennial, CO, USA).

Proteasome inhibitor MG-132 (ab141003) was purchased from Abcam (Cambridge, UK). The eukaryotic protein synthesis inhibitor cycloheximide (CHX) (C7698) was obtained from Sigma-Aldrich (St. Louis, MO, USA).

### Host Cell Culture

The human retinal pigment epithelial cell line ARPE-19 was obtained from the American Type Culture Collection (ATCC, Manassas, VA, USA). Cells were routinely cultured at 37℃ under 5% CO₂ in DMEM/F-12, supplemented with 10% heat-inactivated fetal bovine serum (FBS; Gibco BRL, Grand Island, NY, USA) and 1% antibiotic-antimycotic solution (Gibco BRL, Grand Island, NY, USA). ARPE-19 cells were passaged every 2–3 days using 0.25% Trypsin-EDTA (Life Technologies, Carlsbad, CA, USA).

### Animals and Parasites

Male BALB/c and C57BL/6 mice were obtained from DaeHan BioLink Co. (Chungcheongbuk-do, Korea), and *Phf20* transgenic (TG) mice were generated. All mice were maintained under specific-pathogen-free conditions and were 8 weeks old at the time of initial immunization. Animal studies were conducted with the approval of the Chungnam National University Animal Ethics Committee (Daejeon, Korea). *T. gondii* tachyzoites (RH strain) were maintained in ARPE-19 cells at 37°C under 5% CO_2_ and passaged every 2–3 days. Transgenic RH strains expressing green fluorescent protein (GFP-RH) or red fluorescent protein (RFP-RH) were kindly provided by Dr. Yoshifumi Nishikawa (Obihiro University of Agriculture and Veterinary Medicine, Japan) and maintained under the same conditions as the RH strain.

### Targeted gene silencing

One day before transfection, ARPE-19 cells were seeded in a 6-well plate with 3 mL of normal growth medium. The medium was removed, and 500 μL of fresh DMEM/F12 was added. siRNA duplexes (TBC1D4, PHF20) were diluted in 250 μL of fresh DMEM/F12 to a final concentration of 5–100 nM. Lipofectamine™ RNAiMAX transfection reagent (Life Technologies Corporation, Carlsbad, CA, USA)was gently mixed before use, then 5 μL was diluted in 250 μL of medium and incubated for 5 minutes at room temperature. The diluted siRNA duplexes and diluted Lipofectamine™ RNAiMAX were combined and incubated for 20 min at room temperature. The mixture was added to each well containing cells, bringing the final volume to 1 mL. The plate was gently mixed by hand, rocking back and forth. After a 6-hour incubation at 37°C, the medium was replaced with normal growth medium and cells were incubated for an additional 48 hours at 37°C. Gene knockdown efficiency was assessed by Western blot or immunocytochemistry.

### Preparation of *T. gondii* ESA

Freshly purified tachyzoites (1 × 10^8^) from the peritoneal cavity were incubated at 37°C for 3 hours with gentle agitation in test tubes containing 1.0 mL of Hank’s Balanced Salt Solution (HBSS) (Gibco BRL, Rockville, MD, USA). The supernatant containing ESA was collected after centrifugation at 6,000g for 5 minutes. Protein concentration was measured using the Bio-Rad DC protein assay (Bio-Rad Laboratories, Hercules, CA, USA) with BSA as the standard, and samples were stored at-70°C until use.

### Glucose uptake with 2-NBDG

Glucose uptake was monitored using the fluorescent glucose analog 2-[N-(7-nitrobenz-2-oxa-1,3-diazol-4-yl)amino]-2-deoxy-D-glucose (2-NBDG) (Sigma-Aldrich, St. Louis, MO, USA). Thirty minutes before the end of the treatment, cells were incubated with the fluorophore at a final concentration of 200 μg/mL in glucose-free medium. Ten random fields per sample were captured using a Zeiss AxioObserver inverted microscope (Zeiss, Oberkochen, Germany) equipped with a Colibri illumination system, a Plan-Neofluar 40× objective, and a numerical aperture of 1.4. The green signal from 2-NBDG was captured using an HMR Axiocan monochrome camera controlled by Axiovision software version 3.2 (Zeiss, Oberkochen, Germany). The illumination system used a 470 nm LED and a Zeiss fluorescence filter 61. Flow cytometry analysis was performed as follows: after labeling and washing, cells were detached from the well using 2.5% trypsin-EDTA and resuspended in FACS buffer (1% BSA in PBS). After two washes, the cells were resuspended in Cell-Based Assay Buffer, placed on ice, and immediately subjected to the acquisition of 10,000 events using a FACScan flow cytometer (BD Biosciences, San Jose, CA, USA). Data analysis was performed using BD FACSDiva™ software (BD Biosciences, San Jose, CA, USA).

### Glucose uptake with glucose assay kit

Cells were seeded in 6-well plates and infected with live *T. gondii* tachyzoites (RFP-RH) at the indicated multiplicities of infection (MOI) for 24 hours, or subjected to PHF20 knockdown or overexpression, followed by TgESA treatment. The culture medium was collected and glucose uptake was measured using a glucose assay kit (Shanghai Rongsheng Biotech, Shanghai, China) according to the manufacturer’s instructions.

### Quantitative Measurement of GLUT4 Translocation to the Plasma Membrane by Flow Cytometry

Cells were cultured in 6-well plates and treated with insulin or *T. gondii* ESA for 1 hour, or infected with *T. gondii* for 24 hours. For each sample, 5 μL of primary anti-GLUT4 antibody was mixed with 1 μL of secondary chicken anti-goat IgG antibody conjugated to AlexaFluor 488 in 500 μL of medium and incubated for 10 minutes at room temperature in the dark. After dissociation with trypsin-EDTA, the cultured cells were incubated with the diluted antibody at 37°C under 5% CO_2_ for 30 minutes in the dark. Cells were fixed by adding 0.5 mL of PBS + 1% PFA to each well, without shaking, and incubated for 20 minutes at room temperature in the dark. The pellet was washed twice with 1 mL PBS and resuspended in 0.4 mL PBS + 1% PFA. Samples were analyzed using a FACScan flow cytometer (BD Biosciences, San Jose, CA, USA), and data were analyzed using BD FACSDiva™ software (BD Biosciences, San Jose, CA, USA).

### Solubilized Membrane Extraction

Cells cultured in 100 mm plates were washed with ice-cold PBS, collected, and pelleted by centrifugation at 13,000 rpm for 5 minutes in 1.5 mL microcentrifuge tubes. The supernatants were discarded, and cell pellets were resuspended in 5 volumes of buffer A (20 mM HEPES-KOH (pH 7.5), 10 mM KCl, 1.5 mM MgCl_2_, 1 mM EDTA, 1 mM EGTA, 1 mM DTT, 0.1 mM PMSF, and other protease inhibitors containing 0.25 M sucrose). The mixture was pipetted several times and incubated on ice for 15 minutes. The sample was centrifuged at 1,000g for 10 minutes, and the supernatant was further centrifuged at 10,000g for 30 minutes. The remaining pellet was resuspended in 20 μL of buffer A containing 0.5% CHAPS and centrifuged at 100,000g for 1 hour. The resulting supernatant was the solubilized membrane fraction.

### Translocation of PHF20

ARPE-19 cells seeded in 24-well plates with glass coverslips were infected with RFP-RH at MOI 5 for 24 hours and fixed with 4% paraformaldehyde overnight at 4°C. ARPE-19 cells seeded in 24-well plates with glass coverslips were infected with RFP-RH at MOI 5 for 24 hours and fixed with 4% paraformaldehyde overnight at 4°C. After three washes with PBST, cells were incubated with Alexa Fluor® 488 goat anti-rabbit IgG secondary antibody. After three washes with PBST, cells were mounted using a DAPI-containing mounting solution (Vector Laboratories, Newark, CA, USA) to stain the nucleus. The samples were imaged using a confocal microscope.

### RNA isolation and RT-PCR

Total RNA was extracted from stimulated cells using Trizol reagent (Invitrogen, Carlsbad, CA, USA), and cDNA was synthesized using the ReverTra Ace RT kit (Toyobo, Osaka, Japan). The primer sequences are listed in Table S1. Amplified products were electrophoresed on a 1.5% agarose gel and visualized using Red Safe (Chembio Diagnostics, Medford, NY, USA). mRNA quantification was performed using a NanoDrop spectrophotometer (Thermo Fisher Scientific, Waltham, MA, USA).

### Western blot analysis

SDS-PAGE and Western blot analysis were performed to assess the expression of various proteins. ARPE-19 cells were cultured in 60 mm plates and serum-starved for 4 hours to eliminate stimulation from serum factors. Cells were then stimulated with RH, ESA, or siRNA as indicated. Proteins were extracted using the PRO-PREP™ Protein Extraction Solution (iNtRON Biotechnology, Seongnam, Korea), incubated on ice for 30 minutes, and then boiled for 10 minutes. Equal amounts of protein were loaded onto SDS-PAGE gels and separated by electrophoresis. The separated proteins were transferred onto polyvinylidene difluoride (PVDF) membranes (Bio-Rad Laboratories, Hercules, CA, USA). The membranes were blocked in 5% skim milk in Tris-buffered saline containing 0.1% Tween 20 (TBST) for 1 hour at room temperature. After washing with TBST, the membranes were incubated with the indicated primary antibodies diluted in 1× blocking solution (BioFACT, Daejeon, Korea) overnight at 4°C. After three washes with TBST, the membranes were incubated with HRP-conjugated anti-rabbit or anti-mouse secondary antibodies (Jackson ImmunoResearch Laboratories, Inc., West Grove, PA, USA) for 2 hours at room temperature. The membranes were washed three times and developed using the ProNA™ ECL Ottimo Western Blot Detection Kit (TransLab, Daejeon, Korea).

### Seahorse Assays

Cellular glycolysis was assessed in real-time using the Seahorse XFe24 Extracellular Flux Analyzer (Seahorse Bioscience Inc., North Billerica, MA, USA) by measuring the extracellular acidification rate (ECAR), according to the manufacturer’s instructions. In brief, PHF20 knockdown, overexpressed, or TgESA-treated ARPE-19 cells were plated in 24-well XFe24 plates (3×10^4^ cells/well; Seahorse Bioscience) and incubated overnight at 37°C. The following day, the medium was replaced with XF assay medium and supplemented with glucose, oligomycin, and 2-DG according to the manufacturer’s recommendations. The following day, the medium was replaced with XF assay medium and supplemented with glucose, oligomycin, and 2-DG according to the manufacturer’s recommendations.

## Statistical Analysis

Statistical analysis was performed using the unpaired Student’s t-test. Data from three independent experiments are presented as the mean ± standard deviation (SD). *P* values less than 0.05 were considered statistically significant.

## Results

### 1. *T. gondii* infection promoted glucose uptake in ARPE-19 cells

To evaluate the effect of *T. gondii* on glucose homeostasis in host cells, we measured the intracellular uptake of the fluorescent glucose analog 2-NBDG (2-[N-(7-Nitrobenz-2-oxa-1,3-diazol-4-yl) amino]-2-deoxyglucose) in ARPE-19 cells using fluorescence microscopy. To minimize basal glucose uptake, the cells were serum-starved for 4 hours before being infected with the *T. gondii* RH strain expressing red fluorescent protein (RFP-RH) at different multiplicities of infection (MOI) for 24 hours. The results showed a significant, MOI-dependent increase in 2-NBDG uptake by host cells following *T. gondii* infection. As shown in Figure 1A-B, low levels of *T. gondii* infection (MOI 1) did not significantly increase glucose uptake, whereas MOI 5 resulted in a more than 7-fold increase in 2-NBDG uptake compared to uninfected cells.

**Figure 1.**
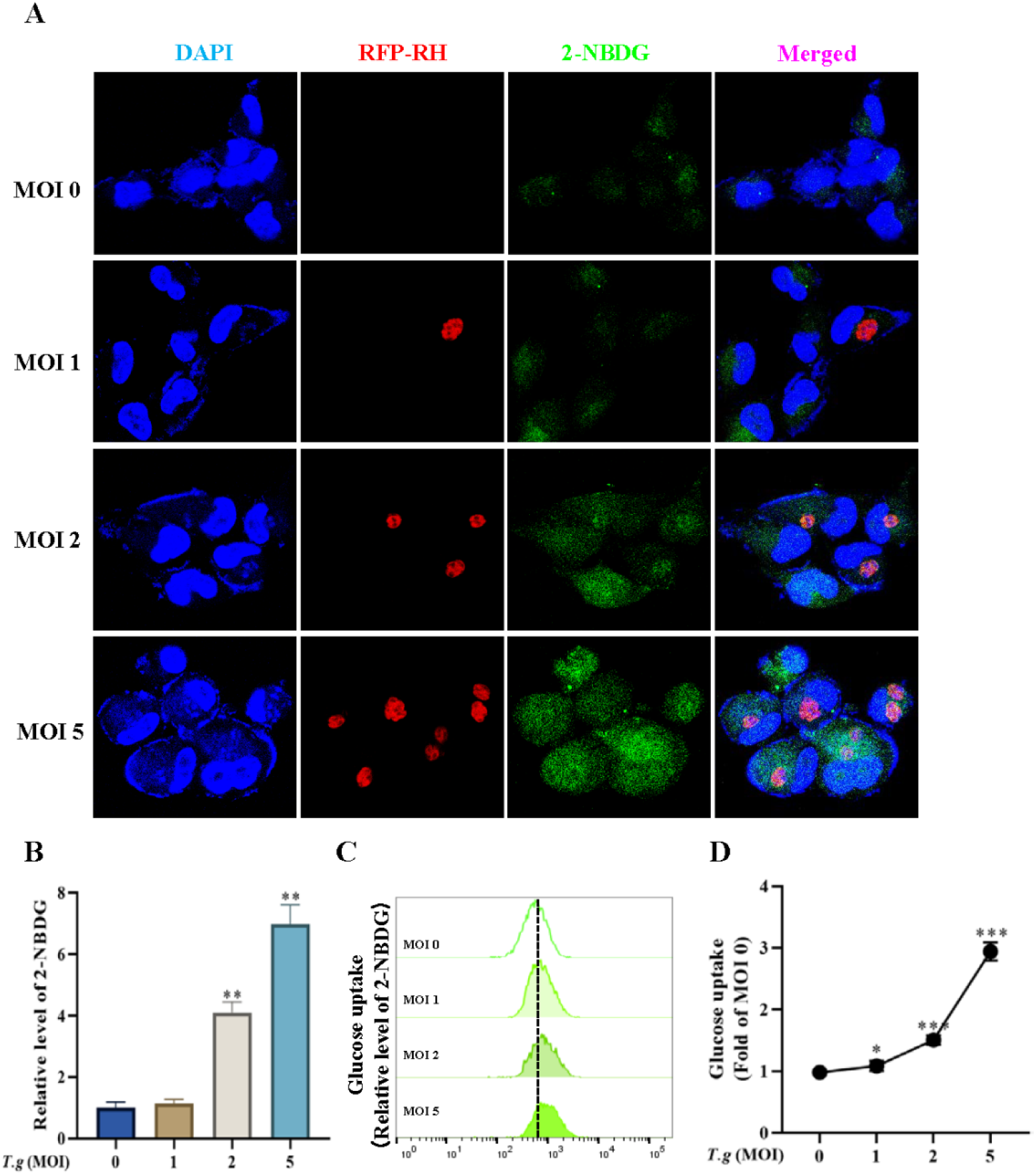
*T. gondii* infection promotes glucose uptake in ARPE-19 cells. (A) Representative fluorescence microscopy images showing the uptake of 2-NBDG (green) in ARPE-19 cells infected with *T. gondii* expressing red fluorescent protein (RFP, red) at different multiplicities of infection (MOI 0, 1, 2, and 5). Nuclei were stained with DAPI (blue). Merged images demonstrate increased 2-NBDG fluorescence in *T. gondii*-infected cells, particularly at higher MOIs. Scale bar, 20 μm. (B) Quantification of 2-NBDG uptake in ARPE-19 cells infected with *T. gondii* at varying MOIs. Data are expressed as the fold change relative to uninfected controls (MOI 0). Results represent the mean ± SD from three independent experiments. ** *P* < 0.01 versus control, as determined by Student’s t-test. (C) Flow cytometry analysis showing a dose-dependent increase in 2-NBDG uptake in ARPE-19 cells with increasing MOI of *T. gondii* infection. The dotted line represents the baseline fluorescence of uninfected cells (MOI 0). (D) Glucose uptake measured using a glucose oxidase-based assay, confirming enhanced glucose uptake in *T. gondii*-infected cells in a MOI-dependent manner. Data are shown as fold change relative to control (MOI 0) with statistical significance indicated: * *P* < 0.05,** *P* < 0.01, *** *P* < 0.001. versus control, as determined by Student’s t-test.

We performed a flow cytometry-based assay to quantify glucose uptake in *T. gondii*-infected host cells using 2-NBDG, a fluorescent glucose analog, and observed a dose-dependent increase in glucose uptake, correlating with the level of infection (Fig. 1C). To further validate these findings, we conducted a glucose oxidase-based colorimetric assay, which confirmed a significant enhancement in glucose consumption following *T. gondii* infection (Fig. 1D). The results showed a dose-dependent increase in intracellular 2-NBDG uptake that correlated with *T. gondii* infection. Consistently, analysis using a glucose oxidase method confirmed that *T. gondii* infection significantly promoted glucose uptake. Taken together, these data suggest that *T. gondii* infection enhances host cell glucose uptake.

### 2. *T. gondii* induced TBC1D4 phosphorylation and also, expression via PI3K/AKT activation

*T. gondii* proliferates within parasitophorous vacuoles, utilizing host glucose to sustain high rates of intracellular replication^[9, 22]^. However, the mechanisms by which the parasite controls host glucose uptake, the impact of glucose on parasite growth, and the functional role of glucose in replication are not fully understood.

To determine whether *T. gondii* manipulates host glucose uptake via TBC1D4 phosphorylation, we measured phospho-TBC1D4 levels by Western blot analysis. Both acute (1 hour) (Fig. 2A) and chronic (24 hours) (Fig. 2B) *T. gondii* infection induced MOI-dependent TBC1D4 phosphorylation. Interestingly, we found that prolonged *T. gondii* infection also increased TBC1D4 protein levels (Fig. 2B). This was reproduced by both live *T. gondii* infection and *T. gondii* secretory/excretory antigen (ESA) treatment (Fig. S1A-B), suggesting that a secretory factor from *T. gondii* is sufficient to activate host glucose uptake.

**Figure 2.**
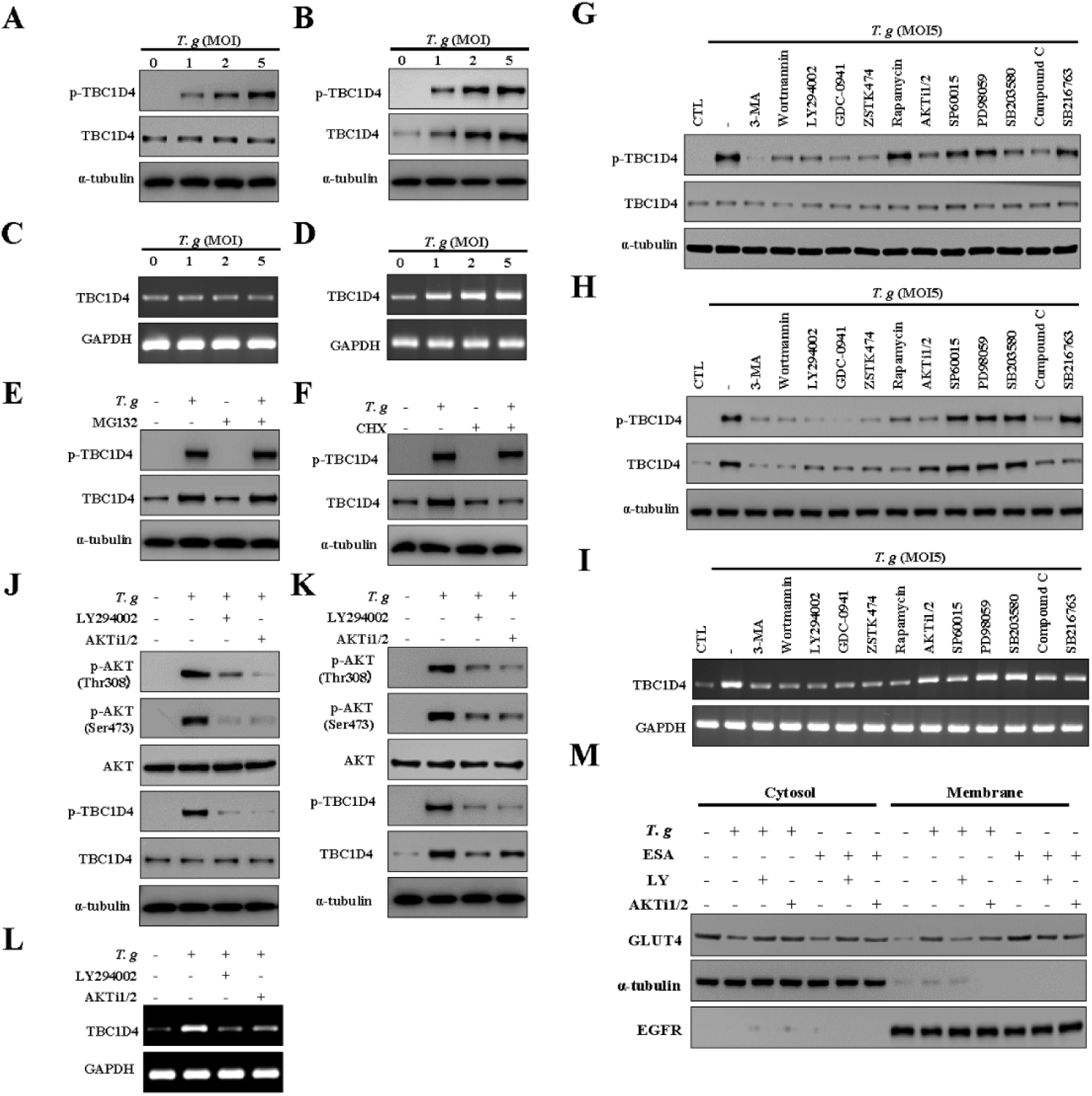
*T. gondii* induces TBC1D4 phosphorylation and expression through the PI3K/AKT signaling pathway. (A, B) Western blot analysis showing the levels of phosphorylated TBC1D4 (p-TBC1D4) and total TBC1D4 in ARPE-19 cells after acute (1 hour, A) and prolonged (24 hours, B) infection with *T. gondii* at varying multiplicities of infection (MOI). TBC1D4 phosphorylation and expression levels increased in a dose-dependent manner. (C, D) RT-PCR analysis demonstrating changes in TBC1D4 mRNA levels following *T. gondii* infection at 1 hour (C) and 24 hours (D). An MOI-dependent increase in mRNA expression was observed at the 24-hour time point. (E, F) The impact of the proteasome inhibitor MG-132 (E) and the protein synthesis inhibitor cycloheximide (CHX) (F) on TBC1D4 protein levels after *T. gondii* infection. MG-132 treatment did not affect TBC1D4 levels, while CHX limited TBC1D4 expression but not phosphorylation. (G-I) Inhibition assays using specific pathway inhibitors to evaluate TBC1D4 phosphorylation and expression at the 24-hour time point. PI3K inhibitors significantly reduced *T. gondii*-induced TBC1D4 activation. (J-L) Western blot analysis of p-AKT (Thr308) and p-AKT(Ser473) levels following *T. gondii* infection with or without the inhibitors of PI3K (LY294002) or AKT (AKTi1/2). Both inhibitors suppressed TBC1D4 phosphorylation. (M) Western blot analysis showing GLUT4 translocation to the plasma membrane in ARPE-19 cells treated with *T. gondii* or excretory/secretory antigen (ESA). The enhancement of GLUT4 translocation was blocked by PI3K/AKT pathway inhibitors, indicating the pathway’s role in mediating this effect.

To further investigate the *T. gondii*-induced changes in TBC1D4 protein levels, we examined whether these changes occurred at the level of mRNA transcription or protein stability. RT-PCR analysis of TBC1D4 mRNA levels after *T. gondii* infection (Fig. 2C) or ESA stimulation (Fig. S1C) showed no significant changes in TBC1D4 gene expression after 1 hour of infection. However, after 24 hours, TBC1D4 mRNA levels increased in an MOI-dependent manner following *T. gondii* infection (Fig. 2D) or in a dose-dependent manner following ESA treatment (Fig. S1D). To investigate the mechanism underlying *T. gondii*-mediated regulation of TBC1D4 protein level, we used the proteasome inhibitor MG-132 and the eukaryotic protein synthesis inhibitor cycloheximide (CHX). Interestingly, MG-132 did not affect TBC1D4 protein abundance after *T. gondii* infection (Fig. 2E) or ESA treatment (Fig. S1E). The role of translation and post-translational modifications in *T. gondii*-induced TBC1D4 regulation should be further investigated. Although CHX limits TBC1D4 levels to basal levels, TBC1D4 is hyperphosphorylated following *T. gondii* infection (Fig. 2F) or ESA treatment (Fig. S1F). These results suggest that *T. gondii* induces TBC1D4 phosphorylation within 1 hour, while prolonged infection or treatment (24 hours) induces both phosphorylation and increased expression of TBC1D4.

TBC1D4 is regulated by PI3K/AKT signaling, prompting us to investigate whether *T. gondii*-induced TBC1D4 phosphorylation is mediated through this pathway. Specific inhibitors of PI3K, mTOR, MAPKs, and other signaling pathways were used to evaluate their effects on TBC1D4 gene expression, protein levels, and phosphorylation at different infection time points. Among these inhibitors, PI3K inhibitors consistently suppressed *T. gondii*-induced (Fig. 2G-I) or ESA-induced (Fig. S1G-I) TBC1D4 phosphorylation and expression. To confirm the involvement of AKT, we used the AKT-specific inhibitor AKTi1/2 and evaluated its effect on TBC1D4. Notably, AKTi1/2 significantly suppressed *T. gondii*-induced (Fig. 2J-L) and ESA-induced (Fig. S1J-L) TBC1D4 phosphorylation and expression. These findings suggest that *T. gondii*-induced TBC1D4 phosphorylation and expression are mediated via the PI3K/AKT signaling pathway.

A key function of insulin is to promote glucose metabolism, primarily through increasing glucose transport. Among the glucose transporters, GLUT4 is predominantly expressed in insulin-responsive tissues, where it mediates glucose uptake upon insulin stimulation. RT-PCR analysis demonstrated that GLUT4 is expressed in ARPE-19 cells, among other glucose transporters (Fig. S2A). Insulin stimulates glucose uptake by triggering the translocation of GLUT4 from intracellular compartments to the cell surface (Fig. S2B). Glucose uptake is regulated by the number of GLUT4 transporters translocated to the plasma membrane. GLUT4 translocation was assessed using flow cytometry (Fig. S2C-D) and Western blot analysis (Fig. 2M) to determine the effect of *T. gondii* on plasma membrane GLUT4 trafficking. *T. gondii* and ESA induced over a 10-fold increase in GLUT4 translocation to the cell surface. This translocation was suppressed by PI3K/AKT inhibitors, indicating the involvement of PI3K/AKT signaling in *T. gondii*-regulated GLUT4 trafficking.

The flow cytometry-based assay showed that *T. gondii*-induced GLUT4 plasma membrane trafficking, along with 2-NBDG uptake, was reversed by the PI3K inhibitor LY294002 (Fig. S2E). This suggests that *T. gondii* facilitates host cell glucose uptake through PI3K/AKT-dependent GLUT4 plasma membrane translocation.

### 3. TBC1D4-dependent glucose uptake is essential for *T. gondii* growth

The Rab GTPase-activating protein TBC1D4, also known as the AKT substrate of 160 kDa (AS160), plays a crucial role in regulating GLUT4 trafficking. While our previous findings indicate that *T. gondii* infection triggers both the phosphorylation and increased expression of TBC1D4 (Fig. 2), its functional role in parasite proliferation remains unclear. To investigate whether TBC1D4 is critical for *T. gondii* replication, we performed fluorescence microscopy to analyze parasite growth in ARPE-19 cells with TBC1D4 knockdown (Fig. 3A). We quantified the *T. gondii* infection ratio by counting infected cells and found that TBC1D4 gene silencing reduced *T. gondii* infectivity (Fig. 3B). Additionally, *T. gondii* proliferation was significantly inhibited by TBC1D4 deficiency. Under normal conditions, more than 45% of GFP-RH tachyzoites replicated three times, producing eight parasites per parasitophorous vacuole (PV), with 2% dividing four times. However, in TBC1D4-knockdown cells, most PVs contained only two or four tachyzoites, and less than 10% replicated three times (Fig. 3C).

**Figure 3.**
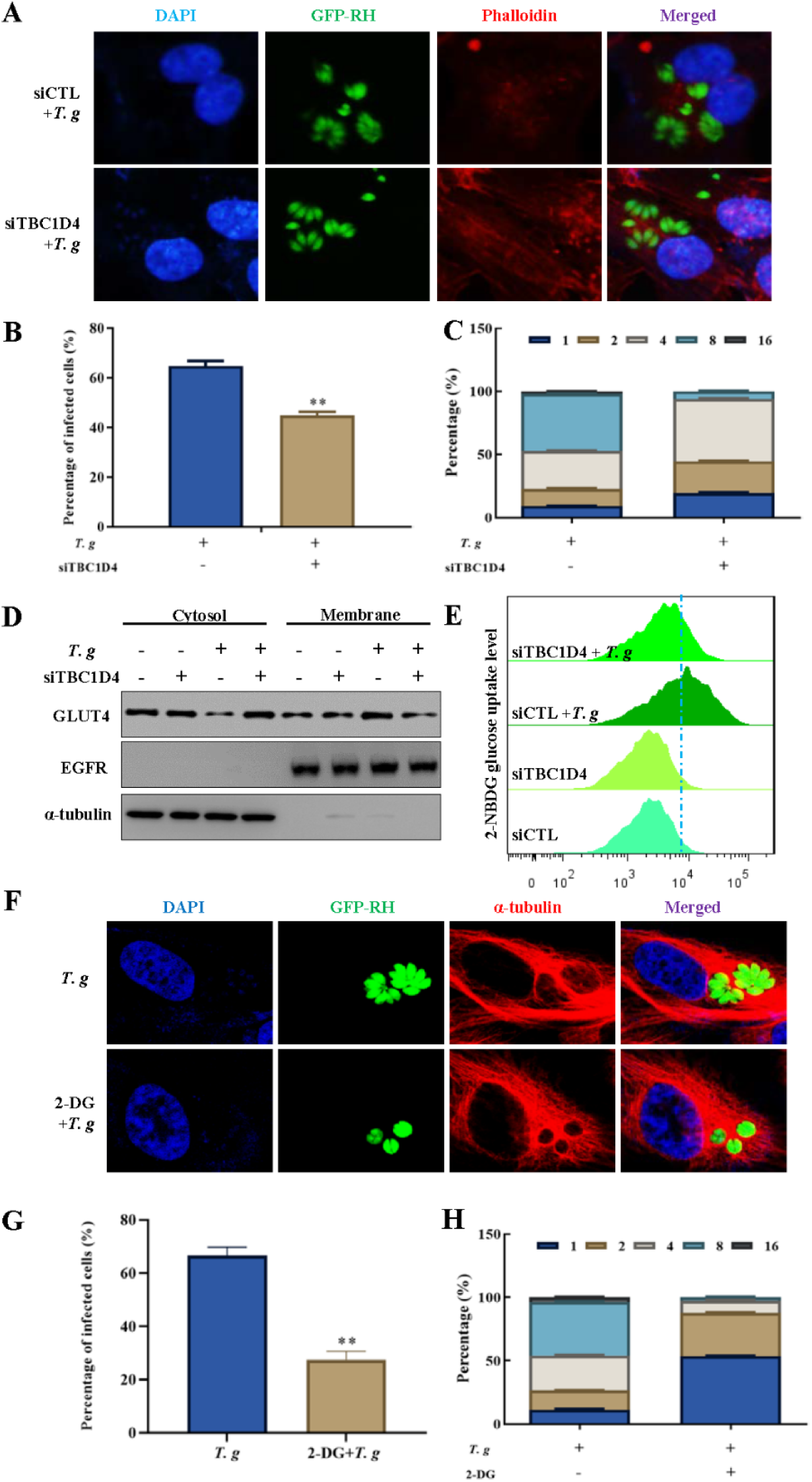
TBC1D4-dependent glucose uptake is essential for *T. gondii* growth. (A) Representative fluorescence microscopy images showing *T. gondii* (GFP-RH, green) replication in ARPE-19 cells, with or without TBC1D4 knockdown. Phalloidin (red) stains the cytoskeleton, and DAPI (blue) marks the nuclei. In TBC1D4-knockdown cells, the replication of *T. gondii* was notably reduced. (B) Quantification of the percentage of infected ARPE-19 cells with or without TBC1D4 silencing. TBC1D4 knockdown significantly reduced *T. gondii* infectivity. Data represent mean ± SD, ** *P* < 0.01. (C) Analysis of *T. gondii* replication, showing the distribution of tachyzoites per parasitophorous vacuole (PV). In TBC1D4-knockdown cells, replication was predominantly limited to two or four parasites per PV, indicating impaired proliferation. (D) Western blot analysis showing the distribution of GLUT4 in the cytosolic and membrane fractions of ARPE-19 cells after *T. gondii* infection, with or without TBC1D4 knockdown. The results indicate reduced GLUT4 translocation in TBC1D4-silenced cells. (E) Flow cytometry analysis of 2-NBDG uptake in ARPE-19 cells under different conditions. TBC1D4 knockdown significantly reduced *T. gondii*-induced glucose uptake. (F) Fluorescence microscopy images showing *T. gondii* replication in cells treated with 2-deoxyglucose (2-DG), which inhibits glycolysis. The replication of *T. gondii* was suppressed in 2-DG-treated cells. (G) The percentage of infected cells after treatment with 2-DG shows significant inhibition of *T. gondii* proliferation. Data represent mean ± SD,** *P* < 0.01. (H) Distribution of tachyzoites per PV in the presence of 2-DG. Most PVs contained fewer parasites compared to untreated controls, indicating that glucose uptake is crucial for *T. gondii* proliferation

TBC1D4 regulates GLUT4-dependent glucose uptake and contributes to cellular energy metabolism (Fig. 3D-E). We investigated whether TBC1D4 deficiency induces autophagy through a metabolism-related pathway. Western blot analysis of microtubule-associated protein 1A/1B light chain 3B (LC3) showed that TBC1D4 depletion did not affect autophagy (Fig. S3C-D). However, *T. gondii* infection in TBC1D4-silenced cells significantly induced autophagy (Fig. S3E). These results identify TBC1D4 as a key link between *T. gondii*-induced PI3K/AKT signaling and the vesicle trafficking machinery responsible for GLUT4 translocation, enhancing host glucose levels and supporting *T. gondii* proliferation.

*T. gondii* utilizes host glucose for energy production and glycoconjugate synthesis, both of which are essential for its survival. To examine the role of GLUT4-mediated host glucose uptake in *T. gondii* growth, 2-deoxyglucose (2-DG), a glucose analog and competitive inhibitor of glucose metabolism, was used to block glycolysis by inhibiting hexokinase. *T. gondii* invasion (1 hour) and proliferation (24 hours) were assessed using flow cytometry (Fig. S3A-B). We found that in the absence of 2-DG, approximately 17% of tachyzoites invaded host cells, while 15% invaded in the presence of 2-DG. This minor inhibition of invasion suggests that other metabolic pathways, in addition to glucose uptake, may provide the energy required for invasion. Notably, *T. gondii* proliferation was significantly suppressed by 2-DG (Fig. 3F-G). In the control group, nearly 73% of cells were *T. gondii*-positive, while in the presence of 2-DG, this number dropped to 32%. To further assess the effect of 2-DG on *T. gondii* growth, we counted the number of tachyzoites per parasitophorous vacuole (PV) in ARPE-19 cells using fluorescence microscopy (Fig. 3F). We measured the *T. gondii* infection ratio by counting infected cells and found that 2-DG significantly inhibited *T. gondii* infectivity, as shown by the percentage of infected cells. Under normal conditions, more than 45% of tachyzoites replicated three times, producing eight parasites per PV, with some dividing four times. However, in 2-DG-treated cells, more than 50% of PVs contained only two or four tachyzoites, and less than 5% replicated three times, indicating that host glucose is essential for *T. gondii* replication (Fig. 3H). Taken together, these results demonstrate that *T. gondii* requires host cell glucose for proliferation and infection.

### 4. PHF20 plays a critical role in *T. gondii* survival by transcriptional modulation of TBC1D4 expression in ARPE-19 cells

PHF20/TZP (Tudor and zinc finger domain-containing protein) elicits immune responses in glioblastoma patients and some controls^[23, 24]^, functions as a transcription factor, and has been identified as a novel substrate in the AKT signaling cascade, with its phosphorylation by AKT playing a key role in tumorigenesis^[25, 26]^.

To identify the specific transcription factor regulating TBC1D4, the TBC1D4 promoter region was analyzed. *In silico* analysis of the 1 kb human TBC1D4 promoter sequence identified four potential PHF20 response elements (PREs), corresponding to the consensus PHF20 transcription factor binding sequence (Fig. S4). To confirm whether intracellular TBC1D4 expression is regulated by PHF20, ARPE-19 cells were transfected with PHF20 siRNA or pEGFP-N1-PHF20. Western blot and RT-PCR analyses revealed that silencing endogenous PHF20 reduced host TBC1D4 levels (Fig. 4A), whereas PHF20 overexpression had the opposite effect (Fig. 4B).

**Figure 4.**
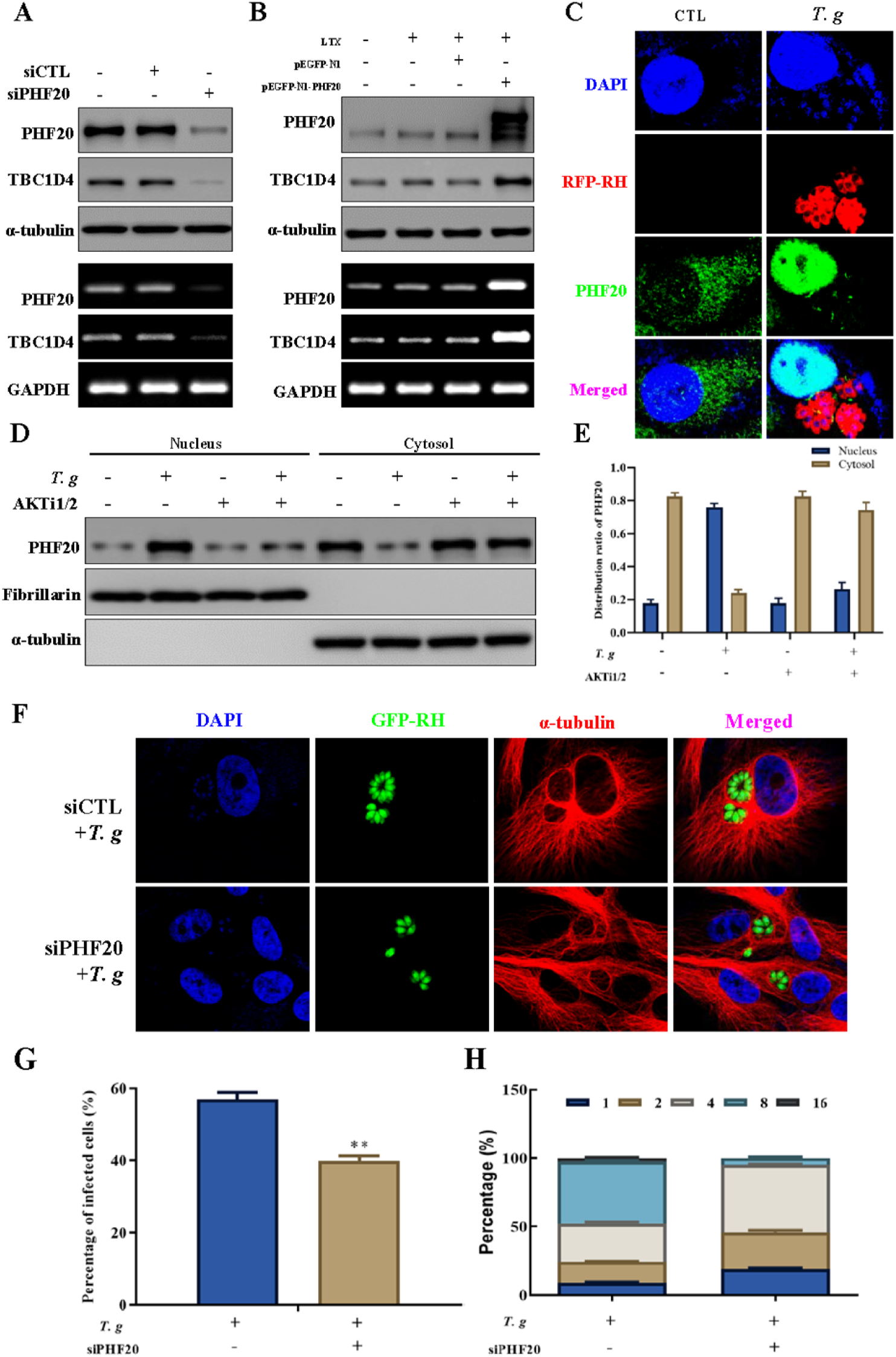
PHF20 is essential for the transcriptional regulation of TBC1D4 expression in ARPE-19 cells during *T. gondii* infection. (A, B) Western blot and RT-PCR analyses showing TBC1D4 and PHF20 expression in ARPE-19 cells transfected with PHF20 siRNA (A) or overexpressing PHF20 (B). Knockdown of PHF20 reduced TBC1D4 levels, while overexpression enhanced TBC1D4 expression. (C) Immunocytochemistry and confocal microscopy showing the localization of PHF20 (green) in ARPE-19 cells, with or without *T. gondii* infection (red, RFP-RH). DAPI (blue) was used to stain the nuclei. During infection, PHF20 translocated from the cytoplasm to the nucleus. (D) Western blot analysis of nuclear and cytosolic fractions demonstrating the redistribution of PHF20 in response to *T. gondii* infection. The majority of PHF20 shifted to the nucleus, a process inhibited by the AKT inhibitor AKTi1/2.(E) Quantification of PHF20 nuclear-to-cytosolic ratio in the presence or absence of AKTi1/2, showing significant nuclear translocation upon *T. gondii* infection. (F) Representative fluorescence microscopy images of *T. gondii* replication in ARPE-19 cells with PHF20 knockdown. The absence of PHF20 significantly impaired *T. gondii* growth. (G) Quantification of the percentage of infected ARPE-19 cells with PHF20 knockdown, showing a significant reduction in *T. gondii* infectivity. Data represent mean ± SD, *P* < 0.01. (H) Analysis of *T. gondii* replication, indicating a decreased number of tachyzoites per parasitophorous vacuole (PV) in PHF20-silenced cells compared to controls.

Immunocytochemistry with confocal microscopy was performed to monitor the intracellular distribution of PHF20. Confocal images showed that during *T. gondii* infection, PHF20 exported from the cytoplasm to the nucleus, where it accumulated (Fig. 4C). Western blot analysis of nuclear and cytosolic fractions was performed to confirm these findings. As shown in Figure 4D, without stimulation, more than 85% of PHF20 was localized in the cytosol. However, nearly 80% of PHF20 translocated to the nucleus following *T. gondii* infection. Nuclear import of PHF20 induced by *T. gondii* was suppressed by the AKT inhibitor AKTi1/2 (Fig. 4E).

Based on these findings, we wondered whether PHF20 is essential for *T. gondii* growth. To test the hypothesis, *T. gondii* replication in PHF20-silenced host cells was monitored using fluorescence microscopy (Fig. 4F). As shown, in the absence of PHF20, *T. gondii* infection (Fig. 4G) and proliferation (Fig. 4H) were significantly suppressed.

Taken together, these data indicate that *T. gondii* infection induces PHF20 nuclear import in an AKT-dependent manner, which is critical for *T. gondii* survival and growth.

### 5. PHF-20 enhanced *T. gondii*-induced glycolysis in ARPE-19 cells

To investigate whether PHF20 is responsible for the uptake of glucose into the cells induced by *T. gondii* infection, we knocked down or overexpressed PHF20 and stimulated cells with ESA to examine its effects on glucose metabolism. PHF20 deficiency reversed ESA-induced glucose uptake (Fig. 5A), whereas PHF20 overexpression enhanced ESA-induced glucose uptake (Fig. 5B).

**Figure 5.**
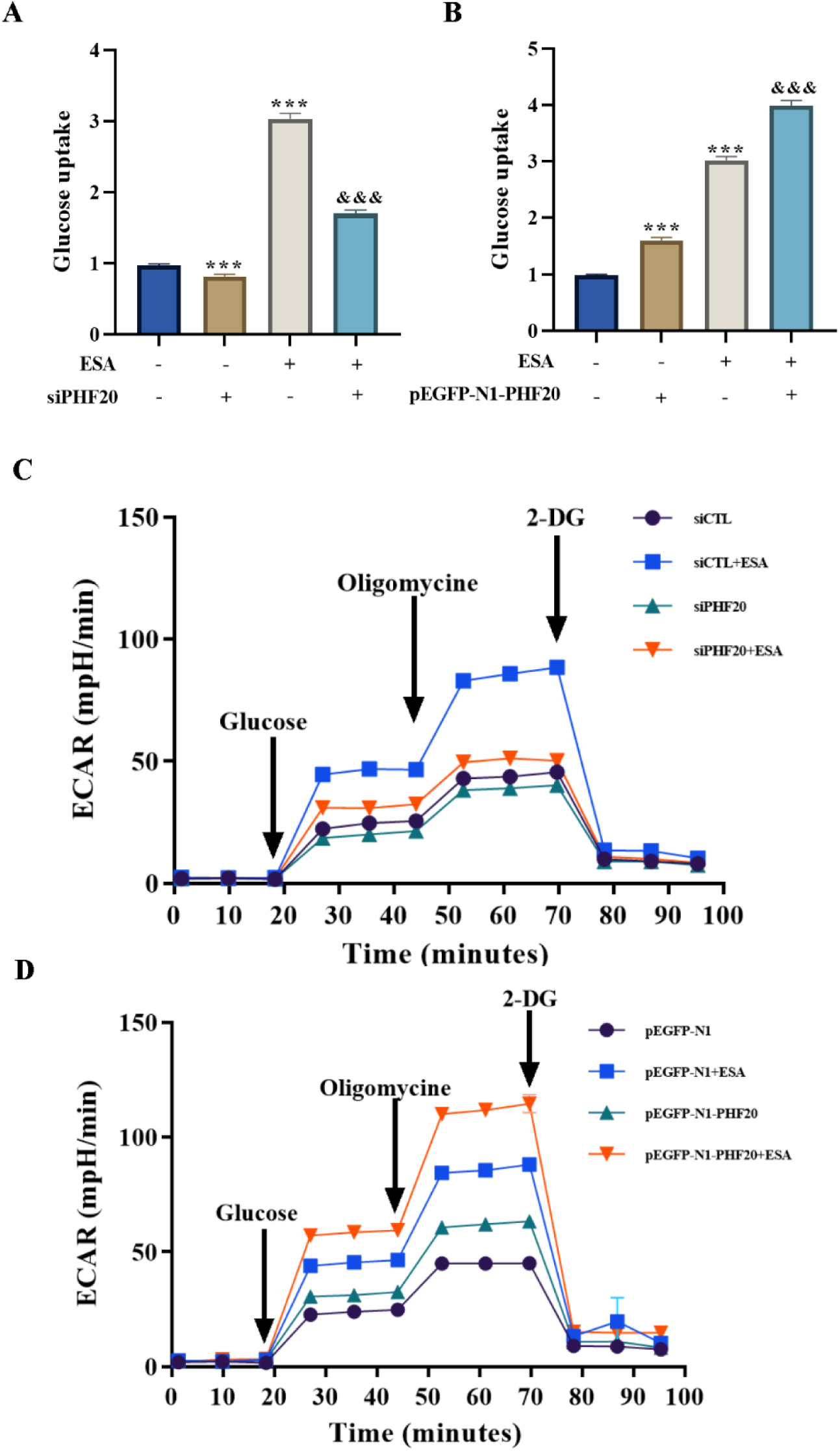
PHF20 enhances *T. gondii*-induced glycolysis in ARPE-19 cells. (A, B) Glucose uptake assays showing the effect of PHF20 knockdown (A) or overexpression (B) on ESA-induced glucose uptake in ARPE-19 cells. Silencing PHF20 significantly reduced glucose uptake, while overexpression enhanced the uptake compared to controls. Data are shown as mean ± SD, *** *P* < *0.001 versus control,* and *&&& P* < 0.001 versus ESA treatment. (C, D) Extracellular acidification rate (ECAR) measurements indicating glycolytic activity in ARPE-19 cells with PHF20 knockdown (C) or overexpression (D) following ESA stimulation. ECAR was measured in real-time following the sequential addition of glucose, oligomycin, and 2-deoxyglucose (2-DG). PHF20 overexpression enhanced ESA-induced glycolysis, while knockdown impaired the glycolytic response.

Intracellular stages of *T. gondii* actively catabolize host glucose via the canonical oxidative tricarboxylic acid (TCA) cycle, a mitochondrial pathway that breaks down organic molecules to generate energy^[27]^. Catabolism of all carbon sources converges at pyruvate, and maintaining a constant pyruvate supply is critical for *T. gondii* growth^[28]^. Glucose in the cell is converted to pyruvate through glycolysis and subsequently to lactate in the cytoplasm. The conversion of glucose to pyruvate, and subsequently lactate, leads to the production and extrusion of protons into the extracellular medium. Proton extrusion acidifies the medium surrounding the cell. The Agilent Seahorse XF Glycolysis Stress Test, which directly measures the extracellular acidification rate (ECAR), was performed to assess the key parameters of glycolytic flux. Results from the ECAR demonstrated that ESA treatment greatly enhanced glycolytic function in ARPE-19 cells, as evidenced by increased glycolysis, glycolytic capacity, glycolytic reserve and non-glycolytic acidification (Fig. 5C-D). Interesting, ESA-induced ECAR was enhanced by PHF20 overexpression (Fig. 5C). However, glycolytic capacity was impaired in PHF20-silenced cells (Fig. 5D). Taken together, our data strongly suggest that PHF20 enhances *T. gondii*-induced glycolysis in ARPE-19 cells.

### 6. PHF20 dependent TBC1D4 regulation is essential for *T. gondii* growth *in vivo*

To validate the previous findings in an *in vivo* model, *Phf20* transgenic mice were used. Eyes (Fig. 6A) and spleens (Fig. 6B) from wild-type and *Phf20* TG mice infected with the RH strain were collected, and parasite burden was assessed using RT-PCR and Western blot analysis. The results indicated that both the eyes (Fig.6A) and spleens (Fig. 6B) of *Phf20* TG mice were more susceptible to *T. gondii* infection compared to wild-type mice.

**Figure 6.**
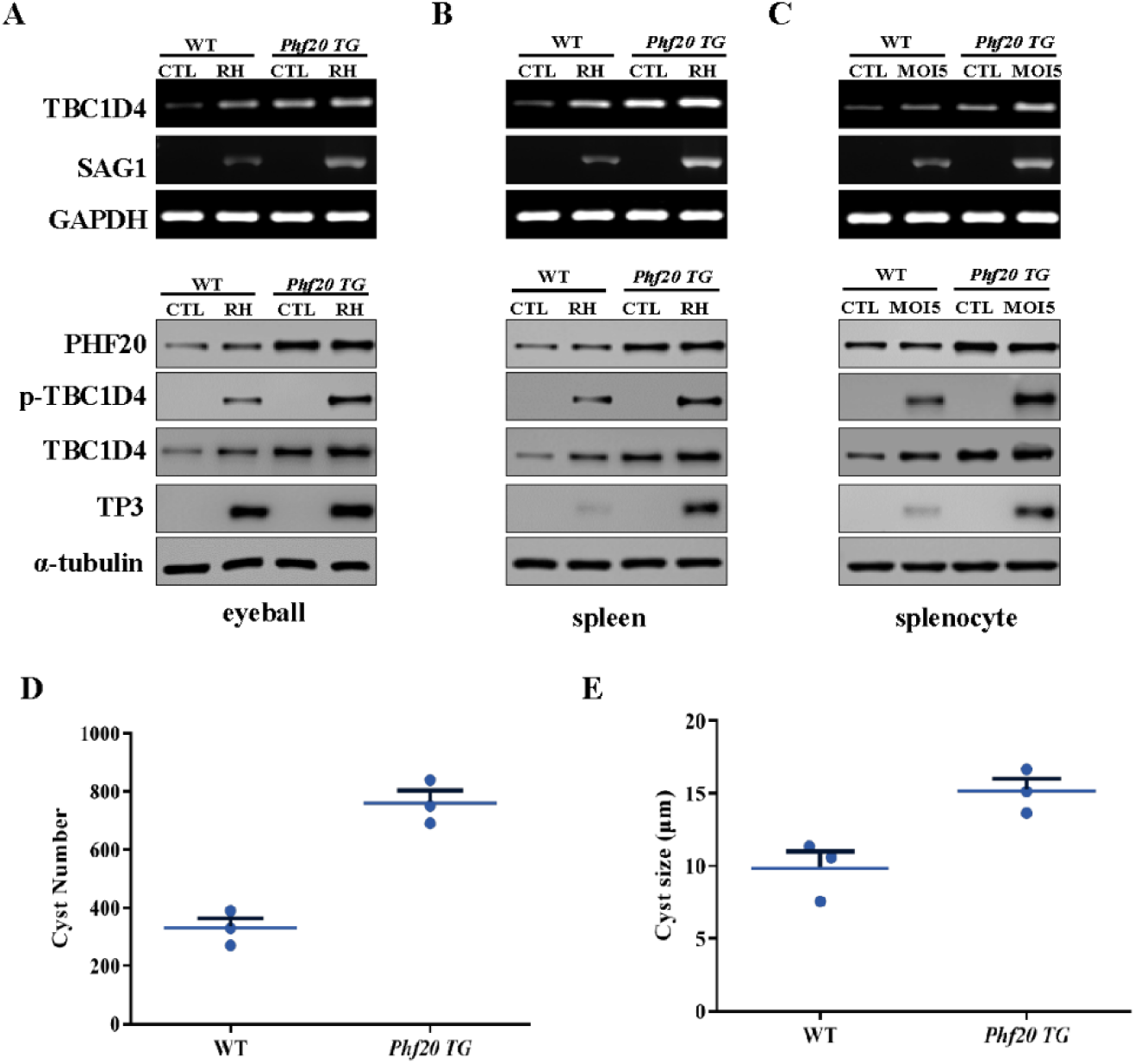
PHF20-dependent TBC1D4 regulation is critical for *T. gondii* growth *in vivo*. (A-C) RT-PCR and Western blot analyses of TBC1D4, PHF20, and *T. gondii* markers (SAG1, TP3) in tissues from wild-type (WT) and *Phf20* transgenic (TG) mice. Samples were collected from the eyeballs (A), spleens (B), and splenocytes (C) following infection with *T. gondii* RH strain or control treatment. *Phf20* TG mice showed higher expression of TBC1D4 and increased susceptibility to infection compared to WT mice. (D, E) Quantification of *T. gondii* cyst burden in the brains of WT and *Phf20* TG mice following oral infection with the ME-49 strain. The number (D) and size (E) of cysts were significantly increased in *Phf20* TG mice compared to WT controls, indicating enhanced susceptibility to infection. Data points represent individual mice, with horizontal lines indicating the mean values.

Consistently, an *ex vivo* infection model using splenocytes from *Phf20* TG mice mirrored biopsy data, showing increased susceptibility of *Phf20* TG splenocytes to *T. gondii* compared to wild-type splenocytes (Fig. 6C).

*T. gondii* bradyzoites reside within a glycan-rich cyst wall, primarily in brain and muscle tissue, and can be transmitted through ingestion of undercooked meat^[29]^. The rapid proliferation of tachyzoites and the slower replication or dormancy of bradyzoites place distinct metabolic demands on *T. gondii*. Proliferation rate and metabolic state are closely interrelated. Bradyzoites undergo substantial remodeling of metabolic homeostasis to cope with nutrient limitations^[30]^. In the mouse model, we investigated whether bradyzoites regulate glucose metabolism via *Phf*20 to support their proliferation. The bradyzoite strain ME-49, frequently associated with human disease, was used to orally infect WT and *Phf20* TG mice, and parasite burden was assessed by counting cysts in brain tissue. The number (Fig. 6D) and size (Fig. 6E) of cysts were significantly larger in *Phf20* TG mice than WT mice.

## Discussion

In this study, we demonstrated a novel strategy of *T. gondii* to take advantage of host PI3K/AKT signaling pathway for the parasite’s survival and proliferation. *T. gondii-*induced TBC1D4 phosphorylation and upregulation promoted GLUT4 membrane trafficking to accelerate host glucose uptake, which is beneficial for the parasite growth (Fig.7).

**Figure 7.**
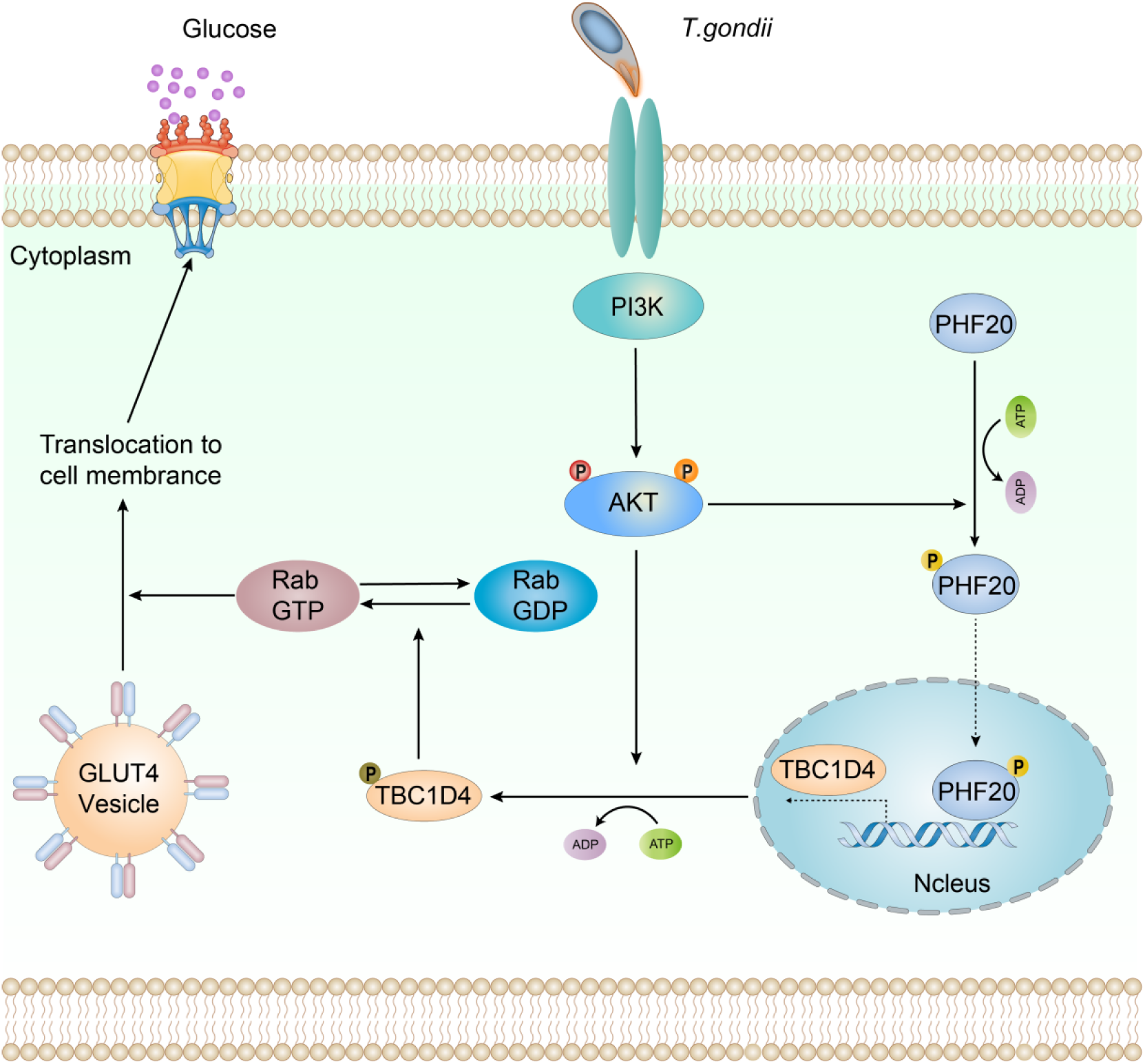
The model of *T. gondii* utilizes host glucose for its growth. Upon infection, *T. gondii* activates PI3K/AKT signaling in host cells, leading to phosphorylation of key downstream effectors. AKT phosphorylates PHF20, promoting its nuclear translocation, enhancing TBC1D4 transcription. Increased TBC1D4 expression and phosphorylation facilitate Rab-GTP to Rab-GDP conversion, which is essential for GLUT4 vesicle translocation to the plasma membrane. This process enhances glucose uptake, ensuring an adequate energy supply to support *T. gondii* intracellular survival and replication.

TBC1D4, also known as AS160, is a Rab GTPase-activating protein that plays a pivotal role in insulin-stimulated glucose uptake by regulating the translocation of the glucose transporter GLUT4 to the plasma membrane. While the phosphorylation and activity of TBC1D4 have been extensively studied, our understanding of its transcriptional regulation is incomplete. It has been reported that TBC1D4 transcript level was increased in T cells from patients with atopic dermatitis, suggesting that its expression can be modulated in certain pathological conditions^[31]^. But still, comprehensive insights into the transcriptional regulation of TBC1D4 under various physiological and pathological conditions remain limited. Further research is needed to elucidate the factors and mechanisms that control the transcriptional dynamics of this critical protein in glucose metabolism.

Here, we identified a multifaceted role of TBC1D4 in facilitating *T. gondii* infection and growth through host cell signaling manipulation. Specifically, we reveal novel insights into the phosphorylation, transcriptional regulation, and functional significance of TBC1D4 during *T. gondii* infection. First, we found that *T. gondii* infection induces TBC1D4 phosphorylation in host cells through the PI3K/AKT signaling pathway, as demonstrated by the inhibition of this modification when PI3K/AKT inhibitors are applied. Previous studies have reported the PI3K/AKT pathway as a key axis manipulated by pathogens to support their survival and replication by regulating host intracellular ROS level^[32, 33]^. These findings highlight the conserved strategy employed by pathogens to hijack host metabolic signaling pathways for their benefit, not only in innate immune response but also in host nutrition regulation.

As a functional consequence of TBC1D4 phosphorylation, *T. gondii-*infection promotes the membrane translocation of GLUT4, one of the well-known glucose transporters. This result is in accordance with previous research indicating that TBC1D4 regulates GLUT4 trafficking and glucose uptake in various cell types, such as HEK293 cells, differentiated 3T3-L1 cells and skeletal muscle cell lines^[34, 35]^. Pathogen-induced GLUT4 translocation has also been reported in other infections, such as *Human Cytomegalovirus* (HCMV), which selectively induces the expression of specific metabolic regulatory kinase, relying on their activity to support glycolytic activation and productive infection^[36]^. This shared mechanism suggests that TBC1D4 and GLUT4 are critical nodes in pathogen-host metabolic crosstalk.

Gene silencing of TBC1D4 significantly impaired *T. gondii* growth and replication, underscoring its essential role in the parasite’s intracellular life cycle. This finding expands the understanding of host-pathogen interactions by highlighting TBC1D4 as a critical host factor for *T. gondii*, similar to studies showing host kinase-dependent survival pathways activated by *T. gondi*^[37, 38]^. Given that glucose is a crucial nutrient for *T. gondii* during intracellular stages^[28]^, our results strongly support that TBC1D4-mediated GLUT4 translocation enhances glucose availability in the parasitophorous vacuole, facilitating optimal parasite growth.

Interestingly, we also discovered that *T. gondii* infection up-regulates TBC1D4 transcription. There was a report that TBC1D4 expression, but not phosphorylation (Ser588, Thr642, Ser704), is critical for elevated insulin-triggered glucose uptake by skeletal muscle from female rats after acute exercise^[39]^, however, the biological mechanism underlying the transcriptional regulation is unexplored. Our signal inhibitor screening confirmed that this transcriptional activation is also mediated by the PI3K/AKT pathway, suggesting a dual role for this signaling in both post-translational and transcriptional regulation of TBC1D4. Although PI3K/AKT has been implicated in transcriptional regulation of various genes^[40, 41]^, the induction of TBC1D4 transcription by this pathway represents a novel mechanism that warrants further exploration. This dual regulation underscores the adaptive capacity of *T. gondii* to ensure sustained availability of host factors critical for its survival.

Plant homeodomain finger protein 20 (PHF20, also termed as glioma-expressed antigen 2) was initially discovered as an autoantibody in patients suffering from glioblastoma^[23, 42]^. Subsequently, it was found that PHF20 was abundantly expressed in various cancers, such as breast cancers^[43]^, colorectal cancers^[44]^, and other diseases associated with renal fibrosis^[45]^, skeletal muscle osteoblastosis and osteoporosis^[46]^. Over the past decade, the functions of PHF20 in several signaling processes have been studied, including those of PKB-mediated phosphorylation^[25]^, autophagy^[47]^, osteogenic differentiation^[48]^ and histone H3 lysine 4 (H3K4) methylation^[49]^, but its role in PHF20 expression refulation has not been reported yet. Our study revealed its biological role in regulating TBC1D4 expression. *In silico* analysis revealed the presence of a conserved binding sequence for PHF20 in the 1 kb human TBC1D4 promoter sequence. PHF20 overexpression increased TBC1D4 levels, whereas siRNA-mediated silencing reduced them. While PHF20 has been identified as a key transcription factor in other contexts^[50, 51]^, its involvement in TBC1D4 regulation provides a new dimension to understand the molecular interplay between host transcription factors and *T. gondii* infection.

Despite these findings, some observations challenge prevailing assumptions. It is well-established that TBC1D4 phosphorylation is generally linked to insulin signaling in metabolic tissues^[52–54]^. Our results reveal a non-canonical activation pathway through *T. gondii* infection, suggesting a broader scope of TBC1D4 function in host-pathogen interactions. This raises the question of whether other pathogens might similarly exploit this pathway. Additionally, while PI3K/AKT signaling is widely recognized for its role in promoting cell survival and metabolism, its involvement in transcriptional up-regulation of TBC1D4, particularly through PHF20, is a less-explored aspect. The potential overlap of pathogen-induced and metabolic signaling pathways could complicate therapeutic targeting, as inhibition of PI3K/AKT may inadvertently affect normal metabolic functions.

Lastly, the observed dependence of *T. gondii* growth on host glucose transport via GLUT4 raises questions about alternative metabolic pathways in parasite-infected cells. Given the emerging evidence of metabolic flexibility in host cells during *T. gondii* infection^[55]^, future studies should explore whether other glucose transporters or metabolic processes compensate for TBC1D4 knockdown.

## Conclusions

In summary, our study uncovers a previously unrecognized host-pathogen interaction in which *Toxoplasma gondii* exploits the PI3K/AKT signaling cascade to induce both phosphorylation and transcriptional upregulation of the host protein TBC1D4. This activation promotes GLUT4 translocation and enhances host glucose uptake, thereby ensuring a steady metabolic supply for parasite proliferation. Notably, we identify PHF20 as a key transcriptional regulator of TBC1D4. AKT signaling promotes the nuclear translocation of PHF20, which in turn is essential for *T. gondii*-induced glycolytic enhancement. Functional assays in both cellular and animal models confirm that PHF20-mediated TBC1D4 expression is critical for optimal parasite growth and persistence. Together, these findings reveal a novel mechanism of metabolic reprogramming during *T. gondii* infection and highlight host glucose metabolism—particularly the PI3K/AKT/PHF20/TBC1D4 axis—as a promising target for therapeutic intervention against toxoplasmosis.

## Declarations

### Ethics approval and consent to participate

All animal experiments were performed in accordance with institutional guidelines and approved by the Institutional Animal Care and Use Committee of Chungnam National University (approval numbers: 202304A-CNU-032 and 202303-CNU-139). Eight-to ten-week-old male C57BL/6 mice were housed under standard conditions with a 12:12-h light–dark cycle, controlled temperature and humidity, and ad libitum access to food and water.

### Consent for publication

Not applicable.

### Availability of data and material

All data generated or analyzed during this study are included in this published article. Further details and supplementary materials are available from the corresponding authors upon reasonable request.

### Authors’ contributions

F. F. Gao, X. Chen, J. Park, and G. H.Cha conceived and designed the study, acquired the data, performed data analysis and interpretation, and drafted the manuscript. F. F.Gao, X.C. Wang, G.H. Hong, Y. S.Yun, W. Zhou, I. W.Choi, and X. Chen performed the experiments. F. F.Gao, X.C. Wang, I. W.Choi, and J. Yuk contributed reagents and materials. X. Chen, J.Park, and G.H. Cha supervised the entire study. All authors discussed the results and approved the final manuscript.

## Acknowledgements

The authors thank Dr. Yoshifumi Nishikawa (Obihiro University of Agriculture and Veterinary Medicine, Japan) for providing GFP-and RFP-expressing *T. gondii* strains.

---

**Figure S1.**
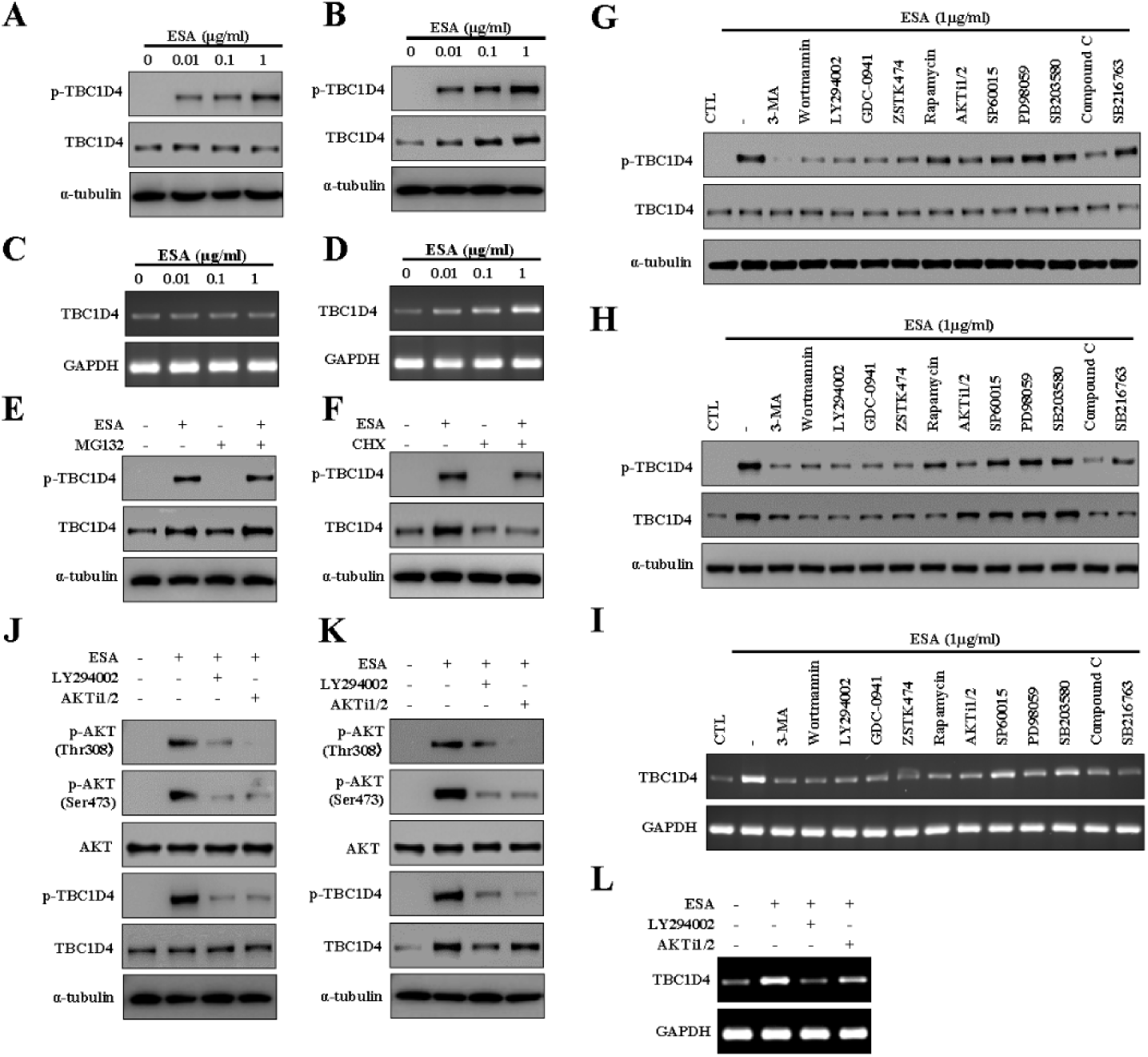
(A, B) Western blot analysis showing dose-dependent phosphorylation of TBC1D4 upon treatment with *T. gondii* excretory-secretory antigen (ESA) at increasing concentrations (0, 0.01, 0.1, and 1 μg/ml) in ARPE-19 cells. Total TBC1D4 and α-tubulin were used as controls. (C, D) RT-PCR analysis of TBC1D4 mRNA levels in ARPE-19 cells treated with ESA at the indicated concentrations. GAPDH was used as a loading control. (E, F) Western blot analysis of TBC1D4 expression in ARPE-19 cells treated with ESA in the presence of MG132 (proteasome inhibitor, E) or cycloheximide (CHX, protein synthesis inhibitor, F). The results indicate that ESA-mediated upregulation of TBC1D4 is primarily regulated at the transcriptional level rather than protein stability. (G-I) Inhibition assay using PI3K, mTOR, and MAPK pathway inhibitors to evaluate the effect of signaling pathway blockade on ESA-induced TBC1D4 phosphorylation (G, H) and transcription (I). Various inhibitors, including LY294002 (PI3K inhibitor), Wortmannin, and Rapamycin (mTOR inhibitors), were used to determine the dependence of ESA-induced TBC1D4 activation on specific pathways. α-tubulin and GAPDH were used as loading controls.(J, K) Western blot analysis confirming that ESA-induced TBC1D4 phosphorylation is dependent on PI3K/AKT signaling. ARPE-19 cells were pre-treated with LY294002 (PI3K inhibitor) or AKTi1/2 (AKT inhibitor) before ESA stimulation. Phosphorylation of AKT at Thr308 and Ser473 was assessed to confirm pathway inhibition.(L) RT-PCR analysis further demonstrating that PI3K/AKT inhibition suppresses ESA-induced TBC1D4 transcription. GAPDH was used as a loading control.

**Figure S2.**
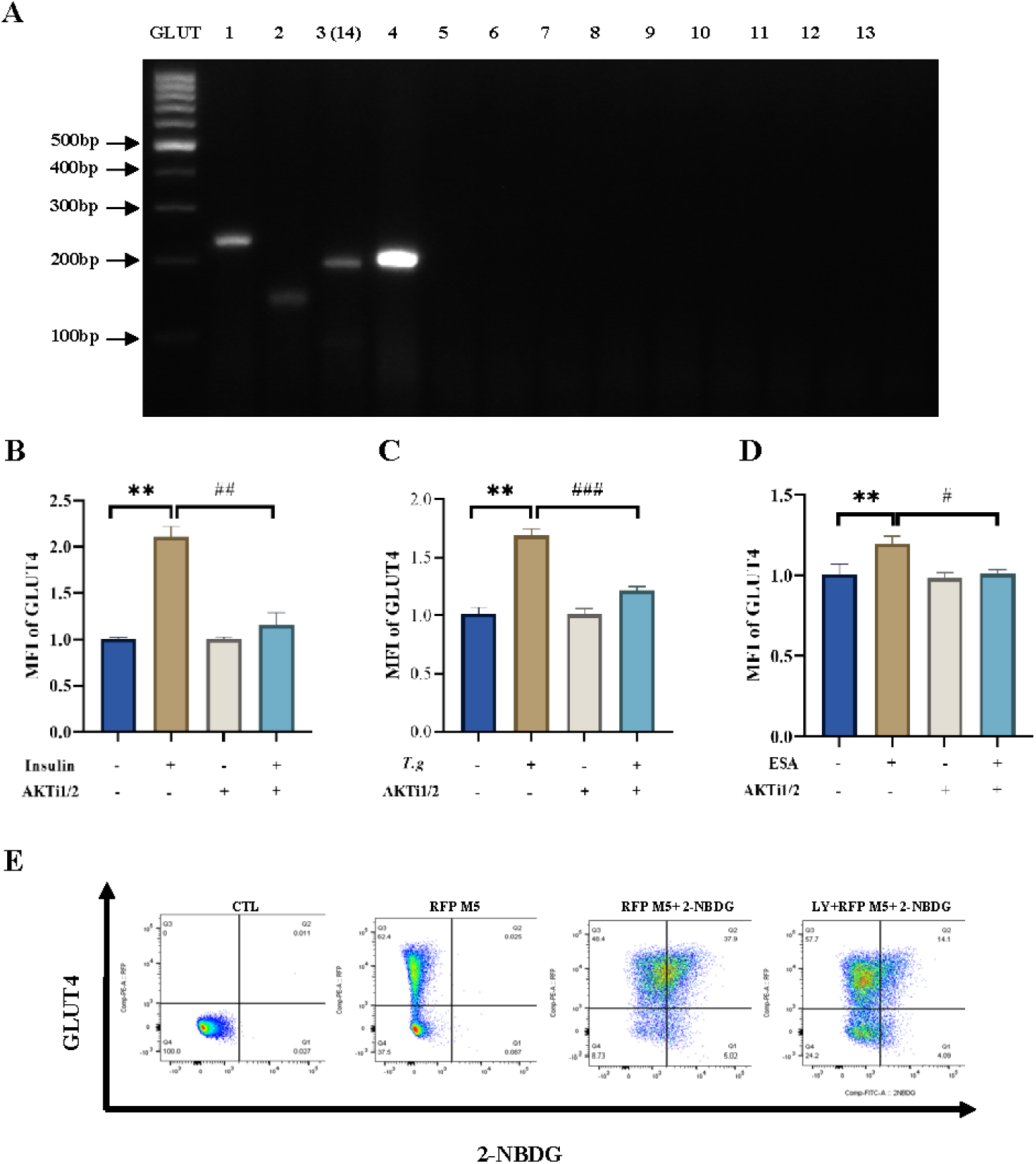
(A) RT-PCR analysis of GLUT4 and other glucose transporters (GLUT) expression in ARPE-19 cells. The gel electrophoresis bands indicate the presence of GLUT4 transcripts, confirming its expression among other GLUT family members.(B-D) Mean fluorescence intensity (MFI) analysis of GLUT4 expression at the plasma membrane under different conditions. (B) Insulin treatment significantly increases GLUT4 MFI, an effect that is suppressed by AKTi1/2, indicating AKT-dependent GLUT4 translocation. (C) *T. gondii* infection enhances GLUT4 translocation, which is also blocked by AKTi1/2, suggesting that *T. gondii*-induced GLUT4 trafficking is mediated via the AKT signaling pathway. (D) ESA stimulation promotes GLUT4 translocation, similar to *T. gondii* infection, and this effect is partially reversed by AKTi1/2.(E) Flow cytometry analysis of GLUT4 translocation and glucose uptake. Representative scatter plots demonstrate increased GLUT4 surface localization in response to *T. gondii* infection and ESA treatment, correlating with enhanced glucose uptake (2-NBDG fluorescence).

**Figure S3.**
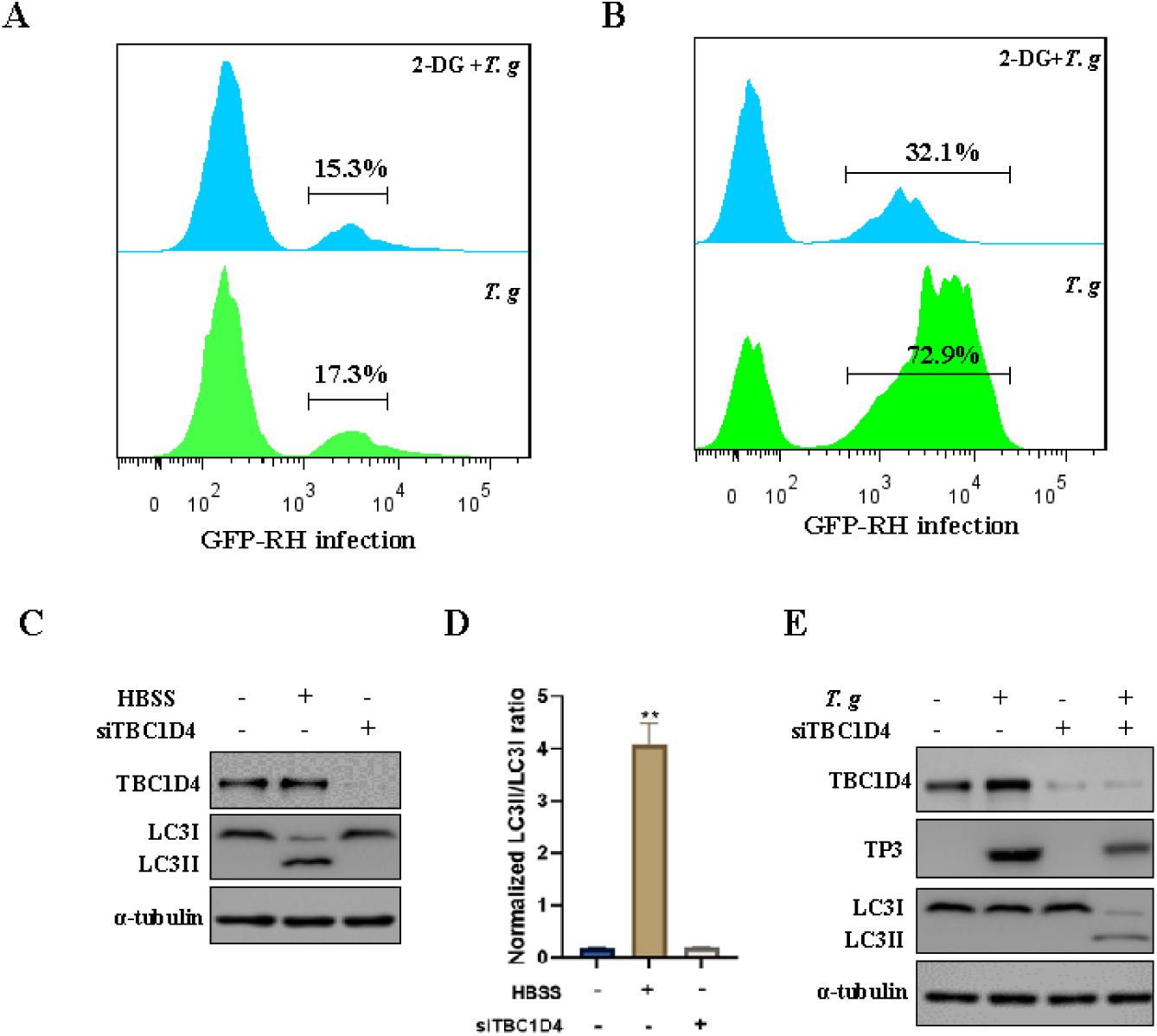
(A, B) Flow cytometry analysis of *T. gondii* infection in host cells treated with 2-DG. Host cells were pretreated with 2-deoxy-D-glucose (2-DG), a glycolysis inhibitor, followed by infection with *T. gondii*-GFP (RH strain). The percentage of infected cells was quantified by flow cytometry. (A) Under normal glucose conditions, *T. gondii* infection rates were comparable between groups, with a slight decrease upon 2-DG treatment. (B) Under high infection conditions, *T. gondii* exhibited significantly higher infection rates, which were substantially reduced by 2-DG treatment (72.9% vs. 32.1%), indicating that host glucose metabolism is crucial for efficient parasite replication. (C, D) TBC1D4 knockdown induces autophagy in host cells.(C) Western blot analysis showing increased LC3-II accumulation in ARPE-19 cells transfected with siTBC1D4, compared to control cells. Cells starved in HBSS (Hank’s Balanced Salt Solution) served as a positive control for autophagy induction. (D) Quantification of the LC3-II/LC3-I ratio, demonstrating that TBC1D4 knockdown significantly promotes autophagy in host cells (*P* < 0.01).(E) TBC1D4 knockdown enhances autophagy during *T. gondii* infection. Western blot analysis showing that *T. gondii* infection increases LC3-II expression in host cells. However, TBC1D4 knockdown further enhances LC3-II accumulation, indicating that loss of TBC1D4 promotes autophagic flux during infection. TP3, a known autophagy regulator, was also upregulated upon TBC1D4 knockdown in infected cells. α-Tubulin was used as a loading control.

**Figure S4.**
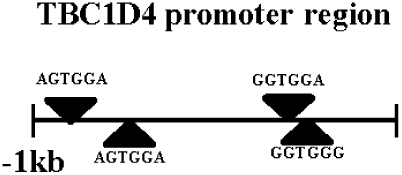
A schematic diagram illustrating the TBC1D4 promoter region (-1 kb upstream of the transcription start site). Putative binding sites for transcription factors are indicated by black triangles. These sites may play a role in the transcriptional regulation of TBC1D4 expression in response to *T. gondii* infection. Sequence motifs identified include AGTGGA and GCTGGC, which are potential regulatory elements involved in metabolic and immune signaling pathways.

**Table S1.**
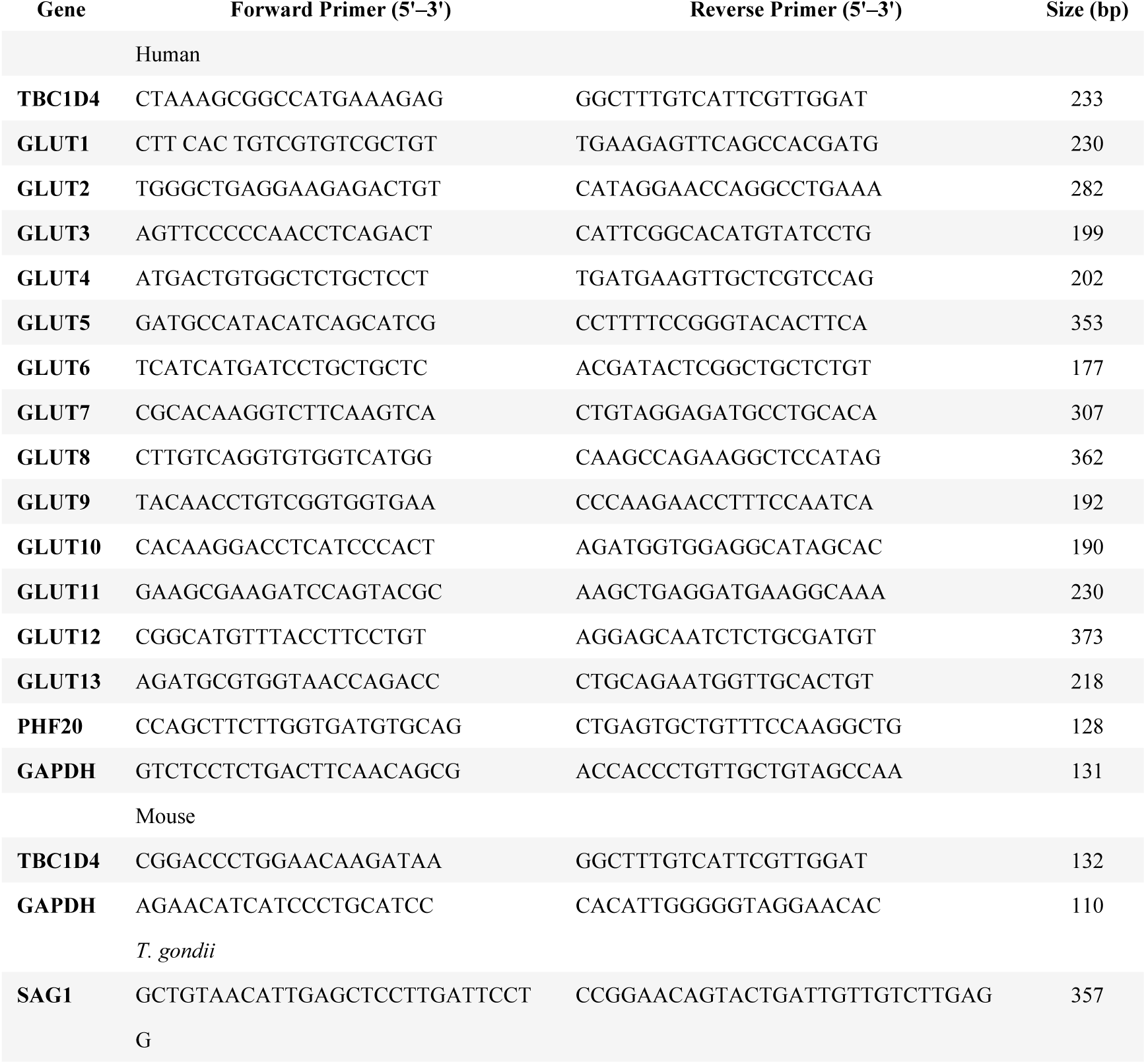
Primers used in this study.

